# Negative Binomial Additive Model for RNA-Seq Data Analysis

**DOI:** 10.1101/599811

**Authors:** Xu Ren, Pei Fen Kuan

## Abstract

High-throughput sequencing experiments followed by differential expression analysis is a widely used approach for detecting genomic biomarkers. A fundamental step in differential expression analysis is to model the association between gene counts and co-variates of interest. Existing models assume linear effect of covariates, which is restrictive and may not be sufficient for some phenotypes. In this paper, we introduce NBAMSeq, a flexible statistical model based on the generalized additive model and allows for information sharing across genes in variance estimation. Specifically, we model the logarithm of mean gene counts as sums of smooth functions with the smoothing parameters and coefficients estimated simultaneously within a nested iterative method. The variance is estimated by the Bayesian shrinkage approach to fully exploit the information across all genes. Based on extensive simulation and case studies of RNA-Seq data, we show that NBAMSeq offers improved performance in detecting nonlinear effect and maintains equivalent performance in detecting linear effect compared to existing methods. Our proposed NBAMSeq is available for download at https://github.com/reese3928/NBAMSeq and in submission to Bioconductor repository.

## 1 Introduction

In recent years, RNA-Seq experiment has become the state-of-the-art method for quantifying mRNAs levels by measuring gene expression digitally in biological samples. An RNA-Seq experiment usually starts with isolating RNA sequences from biological samples using the Illumina Genome Analyzer, the most commonly used platform for generating high-throughput sequencing data. These mRNA sequences are reverse transcribed into cDNA fragments. To reduce the sequencing cost and increase the speed of reading the cDNA fragments (typically a few thousands bp), these fragments are sheared into short reads (50-450bp). These reads are mapped back to the original reference genome and the number of read counts mapping to each gene/transcript region are computed. RNA-Seq experiment is usually summarized as a count table with each row representing a gene/transcript and each column representing a sample.

An important aspect of statistical inference in RNA-Seq data is the differential expression (DE) analysis, which is to perform statistical test on each gene to ascertain whether it is DE or not. Several methods have been developed for DE test using gene counts data, including DESeq2 [1], edgeR [2], which are based on negative binomial regression model and BBSeq [3], which is based on beta-binomial regression model. DESeq2 [1] performs DE analysis in a three-step procedure. The normalization factors for sequencing depth adjustment are first estimated using median-to-ratio method. Both the coefficients and the dispersion parameters in negative binomial distribution are estimated by the Bayesian shrinkage approach to effectively borrow the information across all genes. Finally, the Wald tests or the likelihood ratio tests are performed to identify DE genes. edgeR [2] also uses the negative binomial distribution to model gene counts. It assumes that the variance of gene counts depends on two dispersion parameters, namely the negative binomial dispersion and the quasi-likelihood dispersion. The negative binomial dispersion is estimated by fitting a mean-dispersion trend across all genes whereas the quasi-likelihood dispersion are estimated by Bayesian shrinkage approach [4]. The DE statistical tests are conducted based on either the likelihood ratio tests [5, 6] or the quasi-likelihood F-tests [7] [8]. Both the DESeq2 [1] and edgeR [2] are widely used due in part to their shrinkage estimators which can improve the stability of DE test in a large range of RNA-Seq data or other technologies that generates read counts genomics data. BBSeq [3] is an alternative tool for DE analysis. It assumes that the count data follows a beta-binomial distribution where the beta distribution is parameterized in a way such that the variance accounts for over dispersion. The parameters are estimated by the maximum likelihood approach and DE analysis is based on either the Wald test or the likelihood ratio test. Both the negative binomial and beta-binomial regression models belong to the generalized linear model (GLM) class. GLM is popular owing to the ease of implementation and interpretation, by assuming the link function is a linear combination of covariates. However, as illustrated in the motivating dataset below, the linearity assumption is not always appropriate and nonlinear models for linking the phenotype to gene expression in RNA-Seq data may be important. Stricker et al. [9] proposed GenoGAM, which is a generalized additive model for ChIP-Seq data. However, GenoGAM was developed for a different purpose, i.e., the nonlinear component was to smooth the read count frequencies across genome, and not for relating a nonlinear association between the phenotype and read counts.

In this paper, we introduce NBAMSeq, a generalized additive model for RNA-Seq data. NBAMSeq brings together the negative binomial additive model [10] and information sharing across genes for variance estimation which enhances the power for detecting DE genes, since treating each gene independently suffers from power loss due to the high uncertainty of variance, especially when the sample size is small [1]. By borrowing information across genes, NBAMSeq is able to model the nonlinear association while improving the accuracy of variance estimation and shows significant gain in power for detecting DE genes when the nonlinear relationship exists.

## 2 Motivating Datasets

We demonstrate the existence of nonlinear relationship between gene counts and covariates/phenotypes of interest on two real RNA-Seq datasets. The first dataset is obtained from The Cancer Genome Atlas (TCGA) consortium, a rich repository consisting of high-throughput sequencing omics data for multiple types of cancers; downloadable from the Broad GDAC Firehose. For ease of exposition, we focus on the Glioblastoma multiforme (GBM), an aggressive form of brain cancer. The dataset consists of RNA-Seq count data on 20,501 genes and 158 samples generated using the Illumina HiSeq2000 platform, along with the clinical information of these samples. 16,092 genes were retained after filtering for genes with more than 10 samples having count per million (CPM) less than one. Gene counts are normalized using median-of-ratios as described in DESeq [11] and DESeq2 [1] to account for different library sizes across all samples. Age has been identified as an important prognostic marker for malignant cancers including GBM [12]. To better understand the prognostic value of age in GBM, here we aim to identify age specific gene expression signature. For each individual gene *i*, three models were fitted: (i) base model: *y_ij_* = *β*_0_ + *β*_1_*x*_1*j*_ + *β*_2_*x*_2*j*_, (ii) linear model: *y_ij_* = *β*_0_ + *β*_1_*x*_1*j*_ + *β*_2_*x*_2*j*_ + *β*_3_*x*_3*j*_, (iii) cubic spline regression with 3 knots: *y_ij_* = *β*_0_ + *β*_1_*x*_1*j*_ + *β*_2_*x*_2*j*_ + *f*(*x*_3*j*_), where *y_ij_* denotes logarithm transformed normalized counts of sample *j* and *x*_1*j*_, *x*_2*j*_, *x*_3*j*_ denotes gender, race, age respectively. Race is a binary variable indicating if the sample is Caucasian or non-Caucasian. We then compared the three models using ANOVA F-test, and the p-values are adjusted using the Benjamini &Hochberg [13] false discovery rate (FDR). The test statistics, degrees of freedom of F-statistics, number of significant genes at FDR < 0.05 are summarized in Web Table 1, where SSE_1_, SSE_2_, SSE_3_ denote sum of squared error for model (i), (ii), (iii) respectively. 250 genes are significant when comparing model (i) to model (iii). We further investigated the p-values of these 250 genes when comparing model (i) to model (ii) and found that 30 out of these 250 genes have FDR > 0.2 (see Web Table 2), implying that a simple linear regression is unable to identify these 30 genes even if a less stringent FDR threshold is used. Moreover, 9 genes are significant when comparing model (ii) to model (iii). Figure 1 shows the scatterplots of counts versus age of the most significant genes. For visualization, a local regression is fitted on each scatterplot, suggesting significant nonlinear relationships between normalized gene counts and age. Although only GBM is shown here, we observed similar nonlinear age effect for other types of cancer from Broad GDAC Firehose database (see Web Table 3).

**Figure 1:**
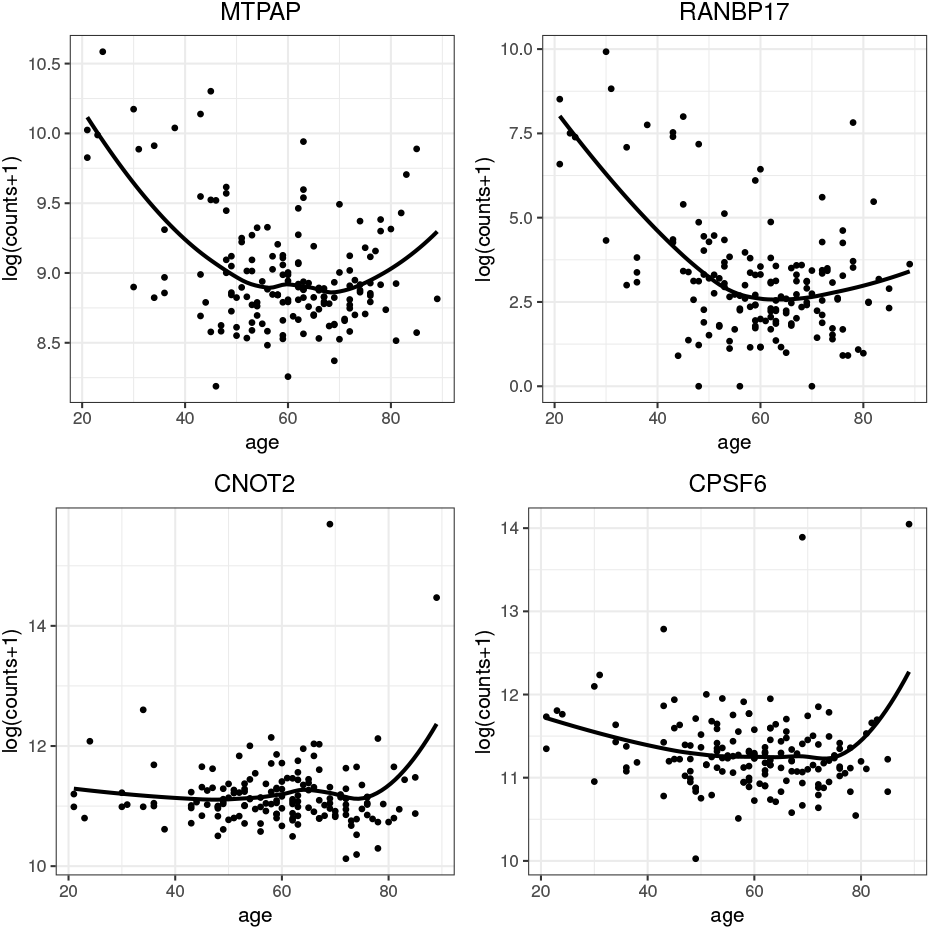
Nonlinear relationship between gene counts and age.

The second dataset comes from our study of gene expression associated with posttraumatic stress disorder (PTSD) in the World Trade Center (WTC) responders [14]. The dataset consists of RNA-Seq read counts of 25,830 genes in whole blood profiled in 324 WTC male responders along with their clinical information. The phenotype of interest is the Posttraumatic Stress Disorder Checklist-Specific Version (PCL), a self-report questionnaire assessing the severity of WTC-related DSM-IV PTSD symptoms [15]. To better understand the nonlinear effect of PTSD severity on gene expression, here we aim to identify gene expression signature related to PCL score. The gene filtering criterion and normalization method are similar to the framework used in the first motivating dataset and samples with missing clinical information were omitted from the analysis. 16,193 genes and 321 samples were retained for DE analysis. In addition, cell type proportions have been implicated in the analysis of whole blood samples. The proportions of CD8T, CD4T, natural killer, B-cell and monocytes were estimated as described in our previous paper [16]. Similar to the first motivating dataset, the base model, linear model, and cubic spline regression model were fitted on each gene to study the effect of PCL score. Age, race, cell proportions of CD8T, CD4T, natural killer, B-cell and monocytes were adjusted in each model. When comparing the cubic spline regression model with base model using ANOVA F-test, 6 genes (AP1G1, NFYA, SCAF11, SPTLC2, TAOK1, ZNF106) are significant at FDR < 0.05. Web Figure 1 shows the scatterplots of counts versus PCL score of these 6 genes with a local regression fitted on each scatterplot. These plots show a consistent pattern, i.e., gene expression first decrease and then gradually increase with the minimum at around PCL score 30.

These observations indicate that the relationship between gene counts and covariate of interest may be nonlinear, and it can be challenging to capture the true underlying nonlinear relationship using parametric methods. Thus, more flexible statistical methods to characterize the nonlinear pattern in RNA-Seq data will improve the DE analysis of complex phenotypes. To this end, we propose a negative binomial additive model to capture the nonlinear association in RNA-Seq data analysis.

## 3 Model Specification

### 3.1 Sequencing Depth Normalization

For RNA-Seq data, the raw counts of each sample are proportional to its sequencing depth i.e. the total number of reads. The raw counts are not directly comparable across samples and therefore normalization factors are needed to adjust for gene counts. Several methods have been developed for RNA-Seq data normalization including total count, upper quartile [17], median, quantile [18, 19], conditional quantile [20], reads per kilobase per million mapped reads (RPKM) [21], median-of-ratios [11], and trimmed mean of M values [22]. A comprehensive comparisons of these normalization methods are provided in Dillies et al. [23] and Li et al. [24]. By comparing the spearman correlation between normalized counts and quantitative reverse transcription polymerase chain reaction (qRT-PCR), Li et al. [24] showed that RPKM yields the best normalization results when alignment accuracy is low. However, by considering the effect of normalization method on DE analysis, Dillies et al. [23] showed that RPKM is ineffective and not recommended. They also showed that the median-of-ratios [11] and trimmed mean of M values [22] maintain low false-positive rate without compromising the power in DE analysis.

Throughout this paper, we adapt the median-of-ratios used by DESeq [11], DEXSeq [25] and DESeq2 [1] for sequencing depth adjustment. Let *K_ij_* be the read count for gene *i* in sample *j*. The normalization factor of sample *j* is defined as:

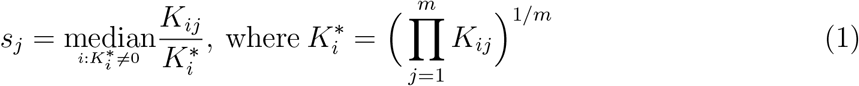

where *m* is the total number of samples.

### 3.2 Generalized Additive Model

For gene *i* and sample *j*, we assume the gene count *K_ij_* follows the generalized additive model:

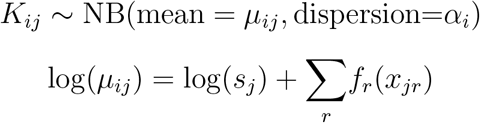

where *μ_ij_* = E(*K_ij_*) is the mean count and *α_i_* is dispersion parameter that relates the mean to the variance by 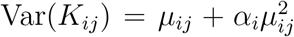, and *x_jr_*’s are the *r* covariates of interest. The logarithm of normalization factor log(*s_j_*) is an offset in the model and *s_j_* is computed by Equation (1) in Section 3.1. *f_r_*(·), *r* = 1, 2,…, *R* represents the spline or smooth function which capture the nonlinear associations between covariate *r* and gene expression. Since our statistical goal is to conduct inference on genes to ascertain whether they are DE with respect to the covariate/phenotype of interest, we choose *f_r_*(·) to be spline functions to facilitate the inference, i.e., 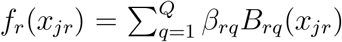, where *β_rq_* are unknown coefficients and *B*(·) are B-spline basis functions. The coefficients ***β*** = (*β*_11_,…, *β*_1*Q*_,…, *β*_*R*1_,…, *β_RQ_*)^*T*^ are estimated by the penalized log-likelihood maximization to avoid overfitting. In particular, the model is estimated by maximizing

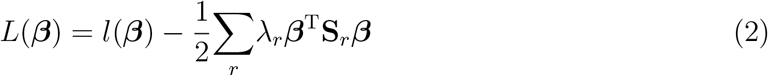

with respect to ***β***, where *l*(·) is the log-likelihood function of negative binomial distribution, λ_*r*_ is smoothing parameter which controls the smoothness of *f_r_*(·), and **S**_*r*_ is a positive semidefinite matrix.

### 3.3 Model Fitting

Several methods have been proposed for fitting the generalized additive model. Hastie and Tibshirani [26, 27] proposed a general backfitting algorithm, whereas Thurston et al [10] adapted the backfitting algorithm for negative binomial distribution. In the backfitting algorithm, given the smoothing parameters λ_*r*_ and dispersion parameters *α_i_*, the coefficients ***β*** can be computed by the penalized iteratively reweighted least squares. However, the most challenging part is in the estimation of the smoothing parameters [28], which is not easy to be incorporated in the backfitting algorithm. Wood et al [29, 30, 31] proposed a nested iterative algorithm which estimates the smoothing parameters and coefficients simultaneously. Their algorithm estimates the smoothing parameters using the cross validation framework in the outer iteration and the coefficients using penalized iteratively reweighted least squares in the inner iteration. We adapted the algorithm implemented in the mgcv package [31, 30, 32, 28, 33] for model fitting, in which various options for model selection criteria and computational strategies are provided. Specifically in NBAMSeq, we adapted the gam function from mgcv [31, 30, 32, 28, 33] for model fitting, in for fitting the generalized additive model on each gene.

The smoothing parameter tuning criteria can be divided into two types, namely (a) the prediction error based criteria including AIC, cross-validation or generalized cross-validation (GCV), and (b) the likelihood based criteria including maximum likelihood or restricted maximum likelihood [30]. The likelihood based methods offer some advantageous over the prediction error based methods since they impose a larger penalty on overfitting and exhibit faster convergence of smoothing parameters estimation [34]. In NBAMSeq, we use the restricted maximum likelihood (REML) method. Maximizing the penalized log-likelihood (2) can be viewed as a Bayesian approach with Gaussian prior 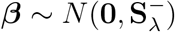, where 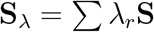 and 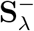 is the inverse or pseudoinverse [35] of **S**_λ_. Under the Bayesian framework, estimating the smoothing parameters can be viewed as maximizing the log-marginal likelihood [29, 30, 31]

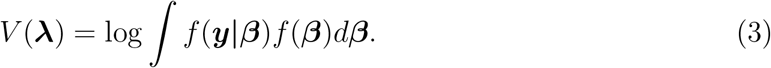

The Laplacian approximation of (3) is given by:

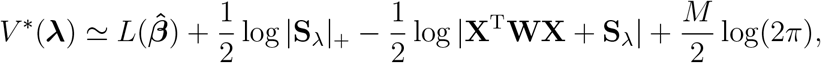

where 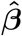 is the maximizer of (2), **X** is the model matrix of covariates i.e. **X** = [**X**^1^, **X**^2^,…, **X**^*R*^] with 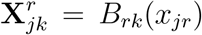. |**S**_λ_|_+_ denotes the product of positive eigenvalues of **S**_λ_, *M* is the number of zero eigenvalues of **S**_λ_, and **W** is the diagonal weight matrix. For negative binomial distribution, the link function is logarithm link and the variance is given by 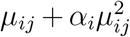, thus the diagonal elements of **W** is given by 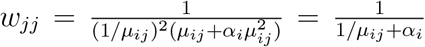. Let λ_*i*1_,…, λ_*iR*_ denote the smoothing parameters of gene *i*. The nested iteration implemented in mgcv [31, 30, 32, 28, 33] can be summarized as Algorithms 1 and 2.

#### Algorithm 1 Outer iteration for **λ** and *α_i_*

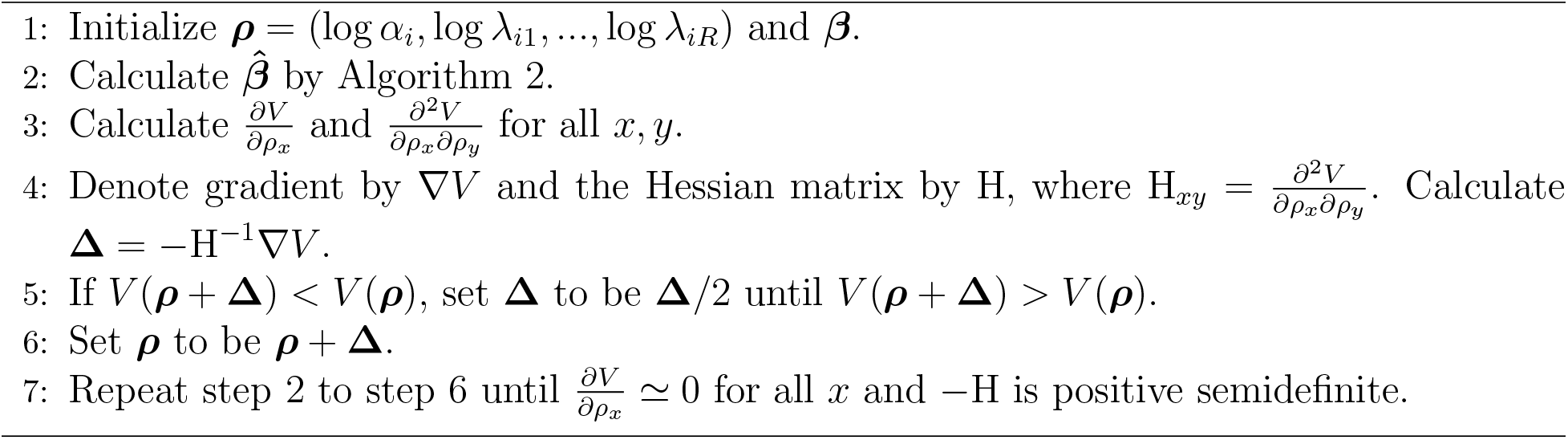

#### Algorithm 2 Inner iteration for β

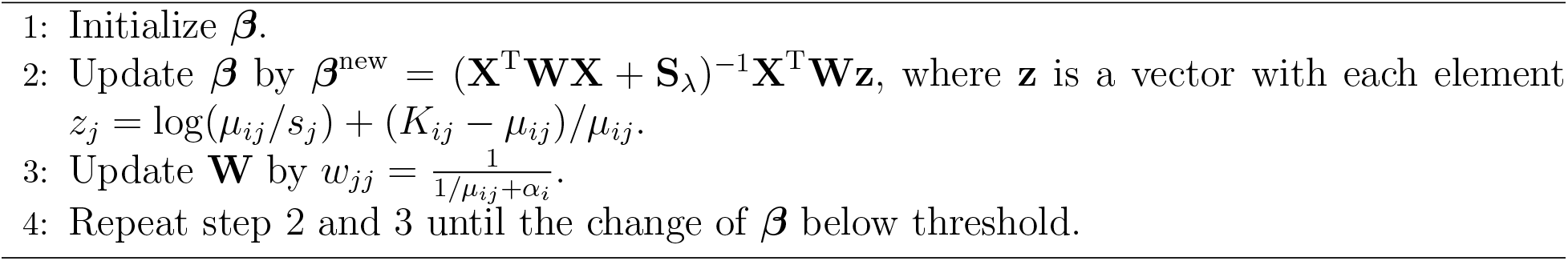

### 3.4 Dispersion Estimation

Information sharing across genes in the estimation of dispersion parameter has been shown to increase accuracy compared to treating each gene independently, especially when the number of replicates is small. Numerous methods have been proposed to incorporate information sharing, and most of them were based on the empirical Bayesian approach which shrinks the gene-wise dispersion estimates toward a common dispersion parameter across all genes.

Following DESeq2 [1], we apply the empirical Bayesian shrinkage dispersion parameter estimate. For each individual gene *i*, we assume a log-normal prior for the dispersion parameters *α_i_*:

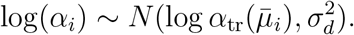

where dispersion trend relates dispersion to mean of normalized 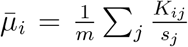 counts by 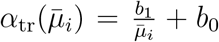 and 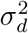 is the prior variance, which is assumed to be constant across all genes. A Gamma regression is fitted to estimate the parameters *b*_1_ and *b*_0_. The prior variance 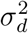 is estimated by subtracting the expected sampling variance of log-dispersion estimator from the variance estimate of residual 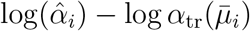 as described in DESeq2 [1]

We first obtain gene-wise dispersion estimates 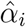 by Algorithms 1 and 2. These gene-wise dispersion estimates are used to fit the dispersion trend and estimate the variance in prior distribution. The final dispersion parameter is estimated by maximizing

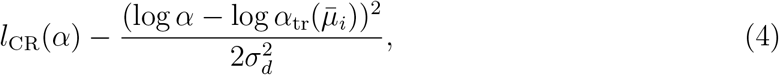

where *l*_CR_ is Cox-Reid adjusted log-likelihood [36]. Note that maximizing (4) can be viewed as a maximize a posterior (MAP) approach. The entire workflow implemented in NBAMSeq is summarized in algorithm 3.

#### Algorithm 3 Workflow of NBAMSeq

1: Estimate normalization factors *s_j_*.

2: Estimate smoothing parameters and gene-wise dispersions by Algorithms 1 and 2.

3: Estimate dispersion trend and prior variance using gene-wise dispersion estimates from Step 2.

4: Calculate MAP estimate of dispersions.

5: Estimate coefficients ***β*** by algorithm 2 using the smoothing parameters in Step 2 and dispersion parameters in Step 4.

## 4 Inference

Our main objective is to identify the genes that exhibit either linear or nonlinear association with the covariate/phenotype of interest. We formulate this as a hypothesis test *H*_0_: *f_r_*(*x_r_*) = 0 versus *H*_1_: *f_r_*(*x_r_*) ≠ 0, where *f_r_*(·) is a smooth function of the covariate of interest. Since we use a Bayesian framework in the coefficients estimation, we have *f*(***β|y***) ∝ *f*(***y|β***)*f*(***β***). In addition, 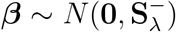, thus 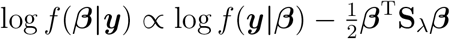. By applying a second order Taylor expansion to 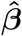, we obtain

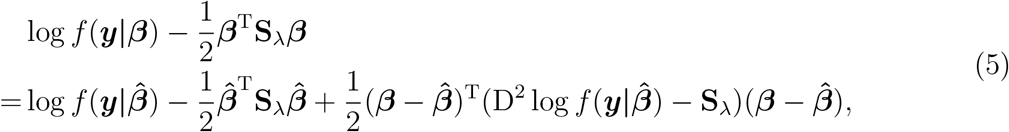

where 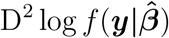 is Hessian matrix of the log-likelihood of negative binomial distribution with respect to ***β***. We use the expected information matrix to replace the Hessian matrix (see Web Appendix A) and equation (5) simplifies to

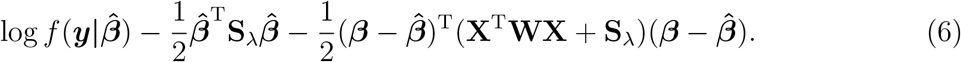

The large sample limit of (6) is the probability density function of multivariate normal distribution, thus 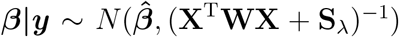 and **V_*β*_** = (**X^T^WX** + **S**_λ_)^−1^ is Bayesian covariance matrix of ***β***. Let **f**_*r*_ be the values of *f_r_*(*x_r_*) evaluated at the observed values of *x_r_* and 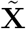 denote the matrix such that 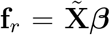. Wood [37] showed that 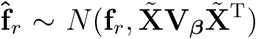 approximately and under the null hypothesis the test statistics 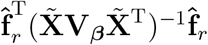 follows a χ^2^ distribution with the degrees of freedom equals to the effective degrees of freedom (edf) corresponding to variable *x_r_* if the edf is an integer. If the edf of *x_r_* is not an integer, Wood [37] showed that the distribution of 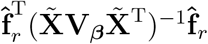 can be approximated by a non-central χ^2^ distribution using the methods of Liu et al [38]. Specifically, the degrees of freedom and the non-central parameter in χ^2^ distribution are determined so that the skewness of 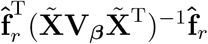 and target χ^2^ distribution are equal, whereas the difference between the kurtosis of 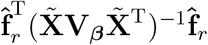 and target χ^2^ distribution is minimized. We utilize this result to calculate a p-value for each gene. The p-values are further adjusted using FDR procedure to account for multiple hypothesis testing.

## 5 Simulation

To evaluate the performance of NBAMSeq, we designed simulation studies in which NBAMSeq was compared with DESeq2 [1] and edgeR [2], two of the most popular softwares for identifying DE genes in RNA-Seq data. Two scenarios were considered, the first was to evaluate the accuracy of dispersion estimates and power of detecting DE genes when the nonlinear effect is true association, the second was to investigate the robustness of NBAMSeq when the linear effect is the true association.

In both scenarios, for gene *i* and sample *j*, we assumed that the gene counts follow negative binomial distribution. i.e. *K_ij_* ∼ NB(mean=*μ_ij_*, dispersion=*α_i_*), *μ_ij_* = *s_j_q_ij_*. For simplicity and without loss of generality, we set all normalization factors *s_j_* to be 1. Since the dispersion parameter decreases with increasing mean counts, we assumed that the true mean dispersion relationship is 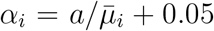. The performance under different degrees of dispersion (*a* = 1, 3, 5) were studied. In practice, although *a* could be larger than 5, the larger value of *a* is compensated by the larger mean of normalized counts. The magnitude of the dispersion *α_i_* is in similar scale regardless of *a*, and thus only *a* = 1, 3, 5 were studied. The covariate of interest *x_j_* was simulated from a uniform distribution U(*u*_1_, *u*_2_). For simplicity, we used *u*_1_ = 20 and *u*_2_ = 80 because 20 to 80 covers the range of age and PCL score in our motivating datasets.

### 5.1 Scenario I

The aim of this scenario was two-fold: (i) to evaluate the accuracy of NBAMSeq dispersion estimates when nonlinear effect is significant, and (ii) to investigate the power and error rate control of NBAMSeq.

We simulated several count matrices which contain the same number of genes (*n* =15,000) across different sample sizes (*m* = 15, 20, 25, 30, 35, 40). We randomly selected 5% of the 15,000 genes to be DE. Within each count matrix, the mean of negative binomial distribution *μ_ij_* was simulated from

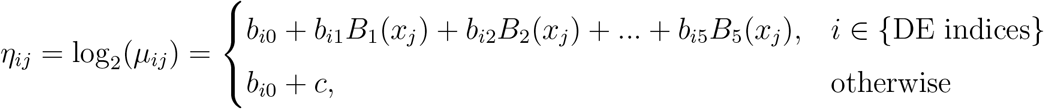

and *μ_ij_* was calculated by *μ_ij_* = 2^*η_ij_*^, where *B*(·) are cubic B-Spline basis functions to induce a nonlinear effect on DE genes, and *c* is a constant to ensure that the count distribution of DE and non-DE genes comparable. The intercepts *b*_*i*0_ were simulated from normal distribution, and the other coefficients *b*_*i*1_, *b*_*i*2_,…, *b*_*i*5_ were simulated from a uniform distribution. The mean and standard deviation of normal distribution, as well as the lower/upper bound of uniform distribution were chosen such that the distribution of 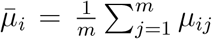 across all genes mimicked our second motivating dataset. Finally, the count of gene *i* in sample *j* was simulated from a negative binomial distribution with mean *μ_ij_* and dispersion *α_i_*. The simulation was repeated 50 times for each combination of dispersion and sample size.

To evaluate the accuracy of dispersion estimates, three methods were compared: (a) DESeq2 [1]; (b) edgeR [2]; (c)NBAMSeq. To illustrate that information sharing across genes produces more accurate dispersion estimates, the gene-wise dispersion estimated by NBAMSeq (NBAMSeq gene-wise) without incorporating prior dispersion trend was also included in the comparison. The average mean squared error (MSE) across 50 repetitions was calculated and the results are shown in Figure 2. Among the DE genes in which the nonlinear effect is significant, NBAMSeq outperforms DESeq2 [1] and edgeR [2] across different sample sizes. On the other hand, among the non-DE genes, the MSE of edgeR [1] is lower than DESeq2 [2] across all degrees of dispersion and achieves comparable performance to NBAMSeq. NBAMSeq also has lower MSE than NBAMSeq (gene-wise), which shows that the Bayesian shrinkage estimation is better than gene-wise dispersion estimation. This simulation scenario shows that NBAMSeq is able to maintain good dispersion accuracy for non-DE genes and shows improvement in the accuracy over DESeq2 [1] and edgeR [2] for DE genes when the nonlinear association is significant.

**Figure 2:**
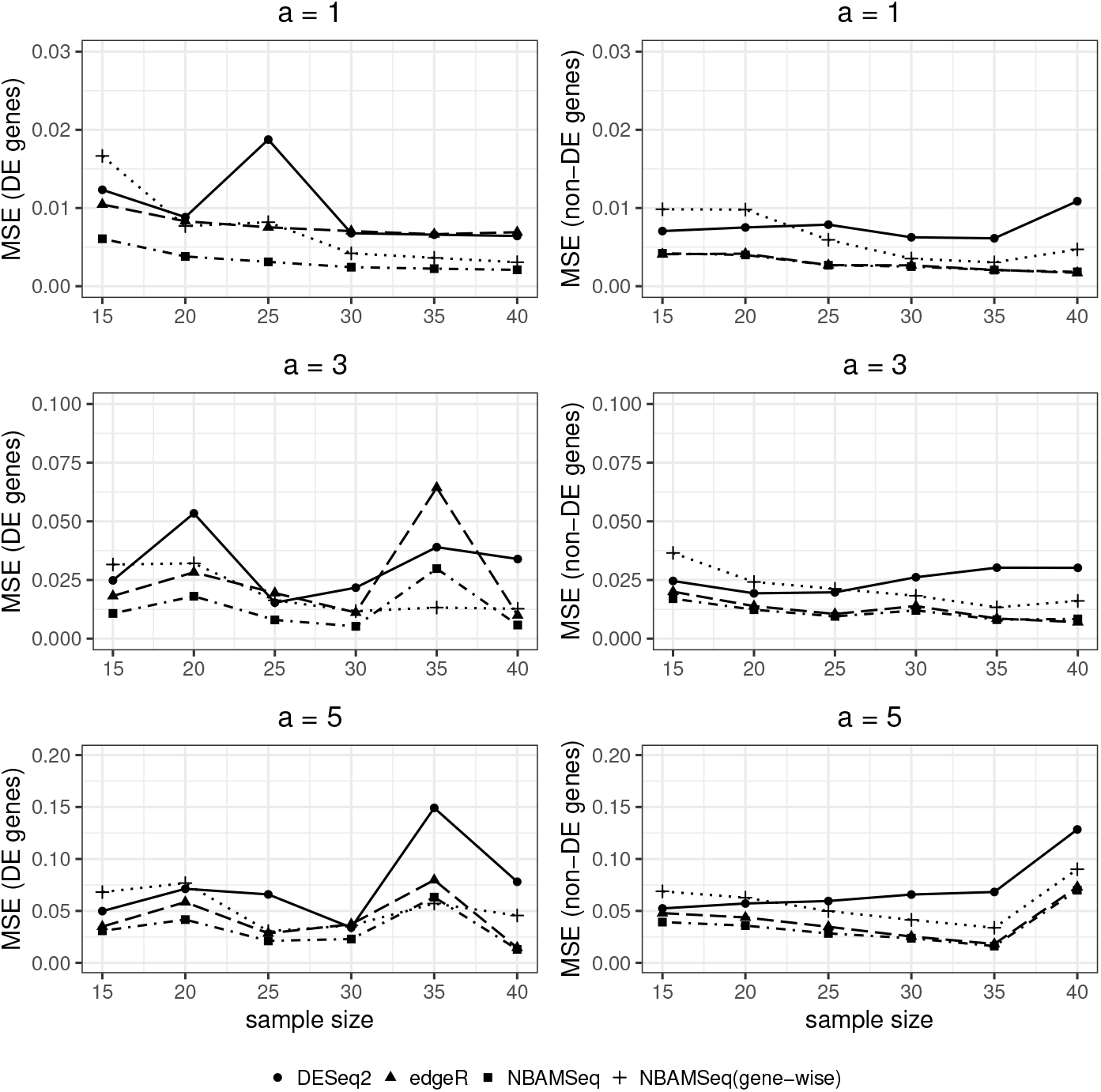
Scenario I: MSE of dispersion estimates.

Figure 3 shows the average true positive rate (TPR), empirical false discovery rate (FDR), empirical false non-discovery rate (FNR) and area under curve (AUC) of DESeq2 [1], edgeR [2] and NBAMSeq. For edgeR [2], two approaches for DE are available, namely the quasilikelihood (QL) and the likelihood ratio test (LRT) approach for calculating the test statistics as described in Lund et al [7]. The authors further showed that detecting DE genes using QL approach controls FDR better than the LRT approach. When calculating TPR, FDR, and FNR, genes are declared to be significant if nominal FDR is < 0.05. The TPR results indicate that our proposed method NBAMSeq has the highest power for detecting nonlinear DE genes regardless of the degrees of dispersion and sample sizes. In addition, NBAMSeq controls the FDR at 0.05 in all cases except in high dispersion (*a* = 5) and low sample size (*m* = 15) in which the FDR is slightly inflated (empirical FDR 0.0576). The FNR results show that NBAMSeq has lower FNR compared to other methods. To ascertain that the power of NBAMSeq is not sensitive to the nominal FDR cutoff for DE genes, the AUCs comparison shows NBAMSeq is consistently more powerful in all cases. The observation also shows that the performance of DESeq2 [1] and edgeR [2] LRT approach are comparable, whereas edgeR [2] QL approach is conservative as shown by the higher FNR. Additional comparisons of the QL approach for NBAMSeq and Gaussian additive models on logarithm transformed count data are provided in Web Figure 2 and 3.

**Figure 3:**
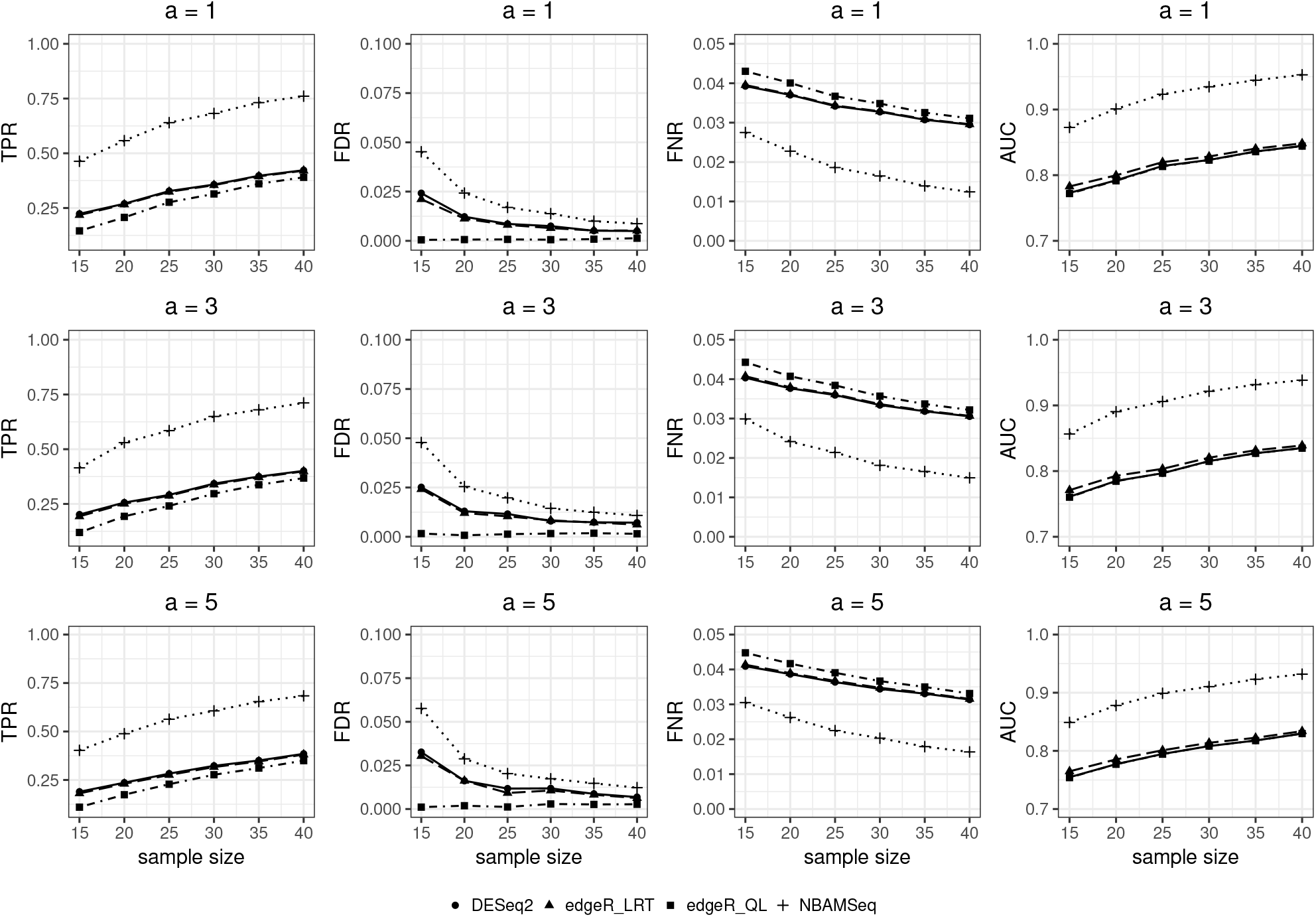
Scenario I: TPR, FDR, FNR, AUC of DESeq2, edgeR and NBAMSeq.

### 5.2 Scenario II

The goal of this scenario was to evaluate the robustness of NBAMSeq when the true association is a linear effect. The simulation setup in this scenario is similar to Scenario I except that *η_ij_* is given by:

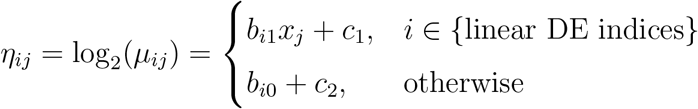

where *c*_1_ and *c*_2_ are constants to ensure that the DE and non-DE genes counts distribution are comparable to avoid bias in detecting DE genes. The linear coefficient *b*_*i*1_ was simulated from normal distribution. Similar to Scenario I, the mean and standard deviation of normal distribution were chosen such that the distribution of 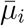 mimicked the distribution of mean normalized counts in real data. The simulation was repeated 50 times and the average MSE of dispersion estimates is shown in Web Figure 4. The accuracy of NBAMSeq is comparable to the accuracy of DESeq2 [1] and edgeR [2] in almost all cases. This simulation shows that although NBAMSeq is designed to detect nonlinear effect, the method is robust and yields accurate dispersion estimates when the true association is linear.

To evaluate the power and error rate control of our proposed model NBAMSeq in detecting DE genes with linear effect, we compare the TPR, FDR, FNR, and AUC at nominal FDR 0.05. Web Figure 5 illustrates that NBAMSeq is a robust approach and maintains comparable performance to other methods.

## 6 Real Data Analysis

In this section, we applied DESeq2 [1] and our proposed method NBAMSeq to our motivating datasets (Section 2), namely the TCGA GBM dataset and WTC dataset. In TCGA GBM dataset, the association between age and each gene expression were investigated, adjusting for sex and race. At FDR 0.05, 845 and 1,439 genes were detected by DESeq2 [1] and our proposed NBAMSeq, respectively. 818 genes were in common between DESeq2 [1] and NBAMSeq, 621 genes were unique to NBAMSeq and 27 genes were unique to DESeq2 [1] (Figure 4). To account for the fact that significant genes are sensitive to the FDR threshold, for the genes unique to NBAMSeq, we investigated their adjusted p-value in DESeq2 [1] (Figure 5A), and vice versa for genes unique to DESeq2 (Figure 5B). A large proportion of the 621 genes unique to NBAMSeq have FDR above 0.1 in DESeq2 [1], implying that DESeq2 [1] is unable to detect the nonlinear genes even if a less stringent FDR threshold is used. However, for the 27 genes unique to DESeq2 [1], their adjusted p-values in NBAMSeq are < 0.1, implying that NBAMSeq is able to detect all these 27 genes at FDR < 0.1. Next, we performed the pathway analysis on DE genes using the hypergeometric tests implemented in the clusterProfiler software [39] for both the Kyoto Encyclopedia of Genes and Genomes (KEGG) [40] and Gene Ontology [41] gene sets. Significant pathways were chosen at FDR 0.05. Gene sets with fewer than 15 genes or greater than 500 genes were omitted from the analysis. For the KEGG pathways, no gene set was significant among DESeq2 DE genes whereas 4 genes sets were significant among NBAMSeq DE genes (Table 1). This result is particularly interesting as mutations in Wnt signaling pathway is implicated in a variety of developmental defects in animals and associated with aging [42, 43]. For the Gene Ontology, a total of 5,000 gene sets were tested, 45 and 78 gene sets were identified by DESeq2 and NBAMSeq, respectively and 34 gene sets were in common between them. A full list of significant gene sets identified by these two models are provided in Web Table 4 and 5. Specifically, among the top gene sets unique to NBAMSeq DE genes (Web Table 6) are extracellular matrix pathways and catabolic process pathways. Changes to extracellular matrix changes with aging has been implicated in the progression of cancer [44], whereas Barzilai et al. [45] showed that metabolic pathways are critical in aging process. These observations suggest that gene sets detected by the NBAMSeq provide additional insights to the development of cancer and aging as a risk factor.

**Figure 4:**
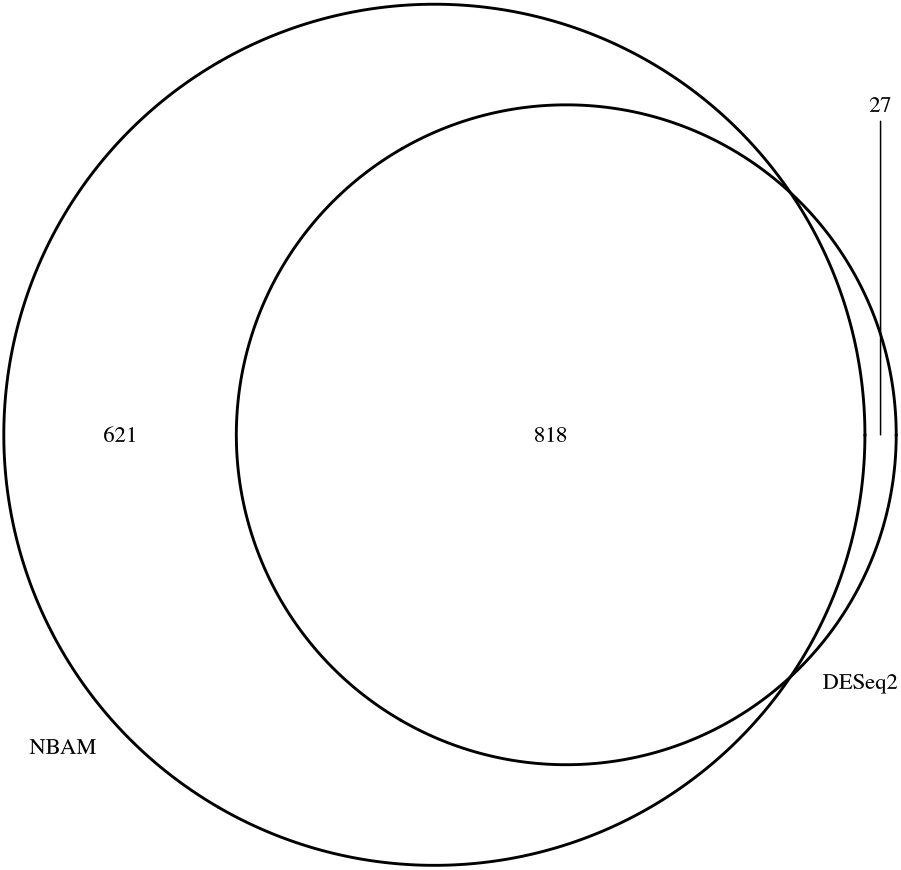
Number of significant genes by DESeq2 and NBAMSeq (GBM dataset).

**Figure 5:**
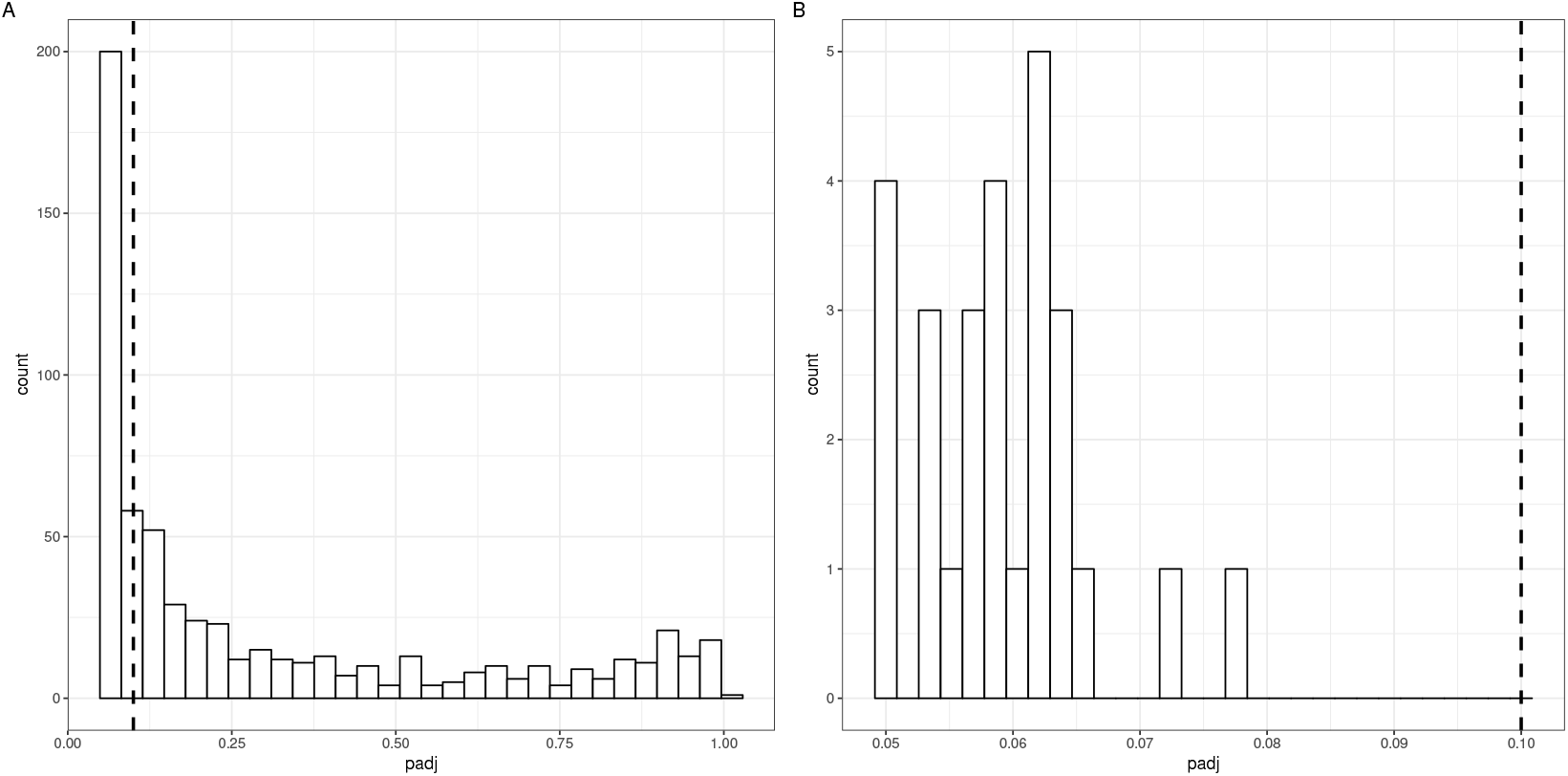
Histogram of adjusted p-values. The blue dashed line is FDR cut-off 0.1.

**Table 1:** KEGG pathways selected by NBAMSeq.

| ID | Description | pvalue | p.adjust |
| --- | --- | --- | --- |
| hsa04310 | Wnt signaling pathway | 0.000252 | 0.033951 |
| hsa04216 | Ferroptosis | 0.000334 | 0.033951 |
| hsa04060 | Cytokine-cytokine receptor interaction | 0.000344 | 0.033951 |
| hsa04974 | Protein digestion and absorption | 0.000463 | 0.034281 |

In the WTC dataset, the relationship between PCL and gene expression were investigated, adjusting for age, race and cell proportions (CD8T, CD4T, natural killer, B-cell, monocytes). At FDR 0.05, 836 and 775 genes were detected by DESeq2 and NBAMSeq, respectively (Web Figure 6). Unlike the TCGA GBM dataset, the genes detected by the two methods have a large overlap, implying that the primary effect of PCL on gene expression is linear. Although NBAMSeq identified fewer PCL gene expression signature compared to DESeq2 [1], the 69 genes unique to DESeq2 were marginal significant in NBAMSeq (Web Figure 7). However, for the 8 genes unique to NBAMSeq, they were not necessarily significant in DESeq2 (Web Table 7). We performed pathway analysis on KEGG [40], and Web Table 8 shows the significant KEGG pathways among NBAMSeq DE genes. As expected, these KEGG pathways were also significant among DESeq2 DE genes due to the large overlapping genes.

## 7 Discussion

This paper introduced a flexible negative binomial additive model combined with Bayesian shrinkage dispersion estimates for RNA-Seq data (NBAMSeq). Motivated by the nonlinear effect of age in TCGA GBM data and PTSD severity (PCL score) in WTC data, our model aimed to detect genes which exhibit nonlinear association with the phenotype of interest. The smoothing parameters and coefficients in NBAMSeq were estimated efficiently by adapting the nested iterative method implemented in mgcv [31, 30, 32, 28, 33]. To increase the accuracy of the dispersion parameter estimation in small sample size scenario, the Bayesian shrinkage approach was applied to model the dispersion trend across all genes. Hypothesis tests to identify DE genes were based on chi-squared approximations.

Although our model was developed for RNA-Seq data, it can be extended to identify nonlinear associations in other genomics biomarkers generated from high throughput sequencing which share similar features as RNA-Seq (e.g., over dispersion in count data). Our simulation studies show that the proposed NBAMSeq is powerful when the underlying true association in nonlinear and robust when the underlying true association in linear compared to DESeq2 [1] and edgeR [2]. In addition, the real data analysis showed that NBAMSeq offers additional biological insights to the role of aging in the development of cancer by modeling the nonlinear age effect. Both the simulation and case studies suggest the advantageous of investigating potential nonlinear associations to detect more effective biomarkers, in order to better understand the biological underpinnings of complex diseases.

NBAMSeq exhibits slight empirical FDR inflation (∼0.06 at nominal FDR 0.05) in small sample size (*n* < 15) and large dispersion scenario, attributed to the chi-squared approximation in the hypothesis tests. One way to circumvent this issue is by using a permutation test to obtain the null distribution of the test statistic, but this approach is computational intensive. Future work includes developing an efficient parallel computing framework for the permutation test. In addition, similar to other softwares for RNA-Seq DE analysis, currently NBAMSeq is developed for inference at gene level in the absence of prior information. Recent methods which take into account the gene network structure in DE analysis have been shown to a powerful approach [46]. Thus, an interesting future research direction is to incorporate the network topology in NBAMSeq to detect nonlinear association and decipher the collective dynamics and regulatory effects of gene expression. Finally, NBAMSeq is available for download at https://github.com/reese3928/NBAMSeq and in submission to Bioconductor repository.

## Acknowledgements

This work was supported in part by the CDC/NIOSH award U01 OH011478.

